# Exploiting Family History in Aggregation Unit-based Genetic Association Tests

**DOI:** 10.1101/2021.04.05.438533

**Authors:** Yanbing Wang, Han Chen, Gina Marie Peloso, Anita DeStefano, Josée Dupuis

## Abstract

The development of sequencing technology calls for new powerful methods to detect disease associations and lower the cost of sequencing studies. Family history (FH) contains information on disease status of relatives, adding valuable information about the probands’ health problems and risk of diseases. Incorporating data from FH is a cost-effective way to improve statistical evidence in genetic studies, and moreover, overcomes limitations in study designs with insufficient cases or missing genotype information for association analysis. We proposed family history aggregation unit-based test (FHAT) and optimal FHAT (FHAT-O) to exploit available FH for rare variant association analysis. Moreover, we extended liability threshold model of case-control status and FH (LT-FH) method in aggregated unit-based methods and compared that with FHAT and FHAT-O. The computational efficiency and flexibility of the FHAT and FHAT-O were demonstrated through both simulations and applications. We showed that FHAT, FHAT-O and LT-FH method offer reasonable control of the type I error unless case/control ratio is extremely unbalanced, in which case they result in smaller inflation than that observed with conventional methods excluding FH. We also demonstrated that FHAT and FHAT-O are more powerful than LT-FH method and conventional methods in many scenarios. By applying FHAT and FHAT-O to the analysis of all cause dementia and hypertension using the exome sequencing data from the UK Biobank, we showed that our methods can improve significance for known regions. Furthermore, we replicated the previous associations in all cause dementia and hypertension and detected novel regions through the exome-wide analysis.

## Introduction

Genome-wide association studies (GWAS) have identified thousands of genetic variants associated with complex diseases at the genome-wide significance level (P< 5×10^−8^). Most of the variants identified by GWAS are common variants with minor allele frequency (MAF)≥ 1%, and most of these variants display modest effect sizes and can only explain a small portion of the total heritability of complex diseases. Yet, rare variants (MAF< 1%) are of vital importance to uncovering unexplained heritability and discovering novel genes contributing to complex diseases. ^1-3^ Because standard association approaches testing each variant individually are grossly underpowered for rare variants, aggregation unit-based methods that jointly analyze variants have been proposed to improve power to detect rare variant associations. Aggregation unit-based approaches include, among others, the sequence kernel association test (SKAT) ^4^, Burden tests ^5- 7^, SKAT-O ^8^, and aggregated Cauchy association test (ACAT) ^9^. However, power of these methods to identify disease regions can be limited by insufficient number of cases in unascertained cohorts.

In genetic association studies, family history (FH) of disease in relatives is often collected in large population cohorts. FH provides an overview of a phenotype within families. Such information typically includes phenotypes of un-genotyped parents or more distant relatives of probands. FH is related to the genotypes of probands at disease loci based on the Mendelian laws of transmission, and is important in assessing health problems and risk of diseases. ^10-12^ While collecting cases is expensive, incorporating FH information into standard case-control genetic association analyses is a cost-effective way to potentially increase statistical power. ^11, 13-15^ Many study designs have limitations for genetic research of late-onset diseases such as Alzheimer’s disease (AD), because disease cases may be deceased with unavailable genotype data. The standard statistical association tests in younger cohorts with low prevalence of some late-onset diseases are not powerful to identify genetic regions associated to a trait of interest. In contrast, the incorporation of available information of disease status in the form of FH may increase the sample size in cohorts with limited cases or individuals with unavailable genotypes. Genetic association studies using only cases and controls will greatly benefit by incorporating available FH information to detect associations.

FH cannot be directly incorporated in standard genetic association methods, limiting its use in genetic association testing. FH has been included as a covariate to improve disease prediction, ^16^ or used to infer mode of inheritance to construct statistical tests. ^17^ However, there are a few reported methods that allow FH to be exploited in genetic association analysis to improve statistical power to detect disease loci. The method developed by Ghosh *et al*. ^13^ enables the incorporation of as a phenotype into the standard single variant analysis, and the results confirmed that exploiting the information contained in FH substantially boosts power to detect the individual variant at disease loci. Nevertheless, these single variant tests suffer from loss of power to detect rare variant associations. While numerous aggregation unit-based methods to jointly analyze rare variants have been proposed to improve power to detect rare variant associations, aggregation unit-based methods that can directly incorporate FH information are needed.

We developed a new and powerful method of family history aggregation unit-based test (FHAT) that enables the incorporation of FH to enhance the statistical power for rare variant associations. We also developed an optimal unified test FHAT-O to maintain robust power in complex scenarios regardless of directions of genetic effects or the proportion of causal variants. To make the comparison with the recent developed method, liability threshold model of case-control status and FH (LT-FH) ^11^, we proposed a novel way to utilize LT-FH into aggregation unit-based method for rare variant analysis (Supplemental Method). We performed an extensive simulation study to evaluate the type I error and power of FHAT and FHAT-O under various scenarios, and illustrated the methods using a real data example from the UK Biobank. We demonstrated that our methods and the LT-FH method control type I error in a reasonable range of significance levels and much better than SKAT, SKAT-O, Burden test, and ACAT when the disease prevalence is low in probands. With greatly reduced computational cost, FHAT and FHAT-O are more powerful than SKAT-LTFH and SKATO-LTFH when the effect of the variant increases with age. FHAT has greater power than SKAT and ACAT-V in most cases when exploiting additional FH information in relatives, and FHAT-O maintains robust power in various complex scenarios. We conducted the rare variant aggregation unit-based tests using unrelated white participants from the UK Biobank tranche of 200,000 individuals with whole-exome sequencing data for all cause dementia (including AD) and hypertension. Based on analysis of known regions associated with all cause dementia and hypertension, we showed that FHAT and FHAT-O have improved significance for most of the known disease regions after incorporating FH especially for all cause dementia given the low prevalence observed in UK Biobank probands. Through an exome-wide analysis, we identified associations with all cause dementia in six genes (*TREM2, PVR, EFCAB3, EMSY, KLC3*, and *ABCA7*) using a suggestive significance threshold (P< 5.6×10^−5^), where *TREM2* reached a stricter significance threshold (P< 2.8×10^−6^). We also found three genes (*GATA5, FGD5, DDN*) associated with hypertension at our suggestive significance level. With enhanced ability to detect disease susceptibility genetic associations, the new findings enabled by these novel methods will contribute to the understanding of the genetic etiology of complex diseases.

## Material and Methods

### Family History Aggregation unit-based Tests (FHAT)

We propose a novel approach, FHAT, to incorporate FH information in the aggregation unit-based tests. We assume that there are *n* probands with *m* observed variants included in the aggregation unit-based test. When we have FH on the relative of the probands, let 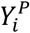 denote the phenotype of the *i*^*th*^ proband; 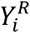 denote the phenotype of the relative of the *i*^*th*^ proband, respectively; 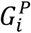 denote the genotypes of the *i*^*th*^ proband; 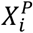 denote covariates for the *i*^*th*^ proband; 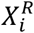 denote covariates of the relative of the *i*^*th*^ proband, such as age and ancestral principal components (PCs) that account for population structure. The probability of observing 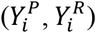 conditional on 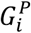 can be written as (see details in Supplemental Method)

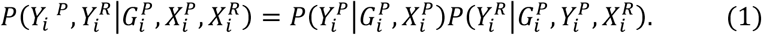

Therefore, the evidence for association can be assessed from two separate analyses for probands and relatives. Based on 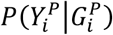, we first assess the association between probands’ genotypes and their disease status using

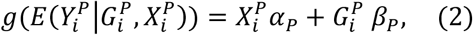

where *g(*·) is the link function, α_p_ is a vector of regression coefficients for covariate effects, β_*p*_ is a vector of regression coefficients for the observed genotypes in probands. The model for relatives based on 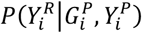 is specified as

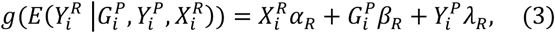

where *λ*_*R*_ is scalar of regression coefficients for probands’ phenotypes for the relatives’ model; *α*_*R*_ is vector of regression coefficients for relatives’ covariates; *β*_*R*_ is vector of regression coefficients for *m* observed variants in probands. This relatives’ model (3) can analyze FH from unrelated relatives, i.e. single relative per probands or FH from both parents since mothers and fathers are conditional independent. We observe that the two underlying association estimators, 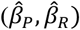, have the relationship ^18^ of 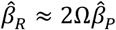, where Ω is the kinship coefficient between probands and their relatives and 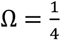 for first-degree relatives such as parents.

Conventional aggregation unit-based methods evaluate the association between a set of variants and phenotype among probands. One such aggregation unit-based method is called the SKAT ^4^. The weighted score statistic based on the probands’ model (2) is

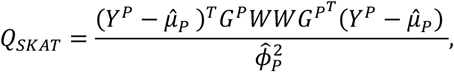

where *W* = *diag(w*_1_, *w*_2_, …, *w*_*m*_) is a pre-specified weight matrix for *m* variants; *G*^*p*^ is a *n × m* genotype matrix with *(i, j*)_*th*_ element corresponding to the additively coded genotype for variant *j* of proband *i*; 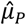 is the estimated mean of *Y*^*p*^using the null model with only covariates; 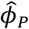 is the dispersion parameter estimate under *H*_*0*_. The score statistic can be obtained similarly to evaluate whether genetic variants are associated with disease status using the relatives’ phenotypes to replace the probands’ phenotypes based on relatives’ model (3). In both probands and relatives analyses, the pre-specified weights can be a function of minor allele frequency. For example, one can use Wu’s weights ^4^ *w*_*j*_ = *Beta (MAF*_*j*_; 1, 2*5*) to up-weight the effect of rarer variants.

We propose to combine the score statistics from the two association models for probands and their relatives using a weighted meta-analysis. Meta-analysis is often used in genetic association analysis to increase the power by combining results from multiple studies. Methods to meta-analyze SKAT results have been developed. ^19^ Meta-analysis of rare variant association tests proposed are based on the study-specific summary statistics, that is, score statistics for each variant and linkage disequilibrium estimates in a region. Because of the genetic relationship between probands and their relatives, we down-weight the scores for relatives by 2Ω when combining the score statistics in a meta-analysis by assuming the homogeneous genetic effects among probands and their relatives. Specifically, because relative *k* of each proband may or may not have phenotype data available, we use 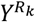 to denote the collective phenotype vector for relative *k* of all probands (e.g., all mothers), including missing values, with kinship coefficient Ω_*k*_. The diagonal matrix *D(R*_*k*_) indicates whether corresponding element in 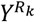 for each proband is missing (denoted by 0) or not (denoted by 1). Therefore, relatives with missing phenotype data do not contribute to the test statistic. We fit a single relative model jointly using all relatives’ phenotypes and covariates conditional on their probands’ phenotypes to get 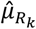, the estimated mean vector of 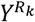 for relative *k* of all probands, as well as the dispersion parameter estimate 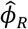 under the null hypothesis of no genetic effects. We assume that all relatives are independent in the model. The general form of FHAT statistics that incorporates FH from relatives is

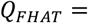

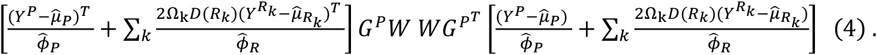

Under the null hypothesis, *Q*_*FHAT*_ follows a weighted sum of chi-square distributions with 1 degree of freedom, 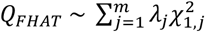. The weights *λ*_*j*_ can be estimated from the eigenvalues of 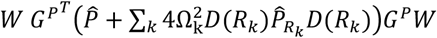, where 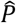 and 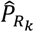 are the projection matrices in probands and relatives *k*, respectively, see Supplemental Method. The p-value can be estimated by the Davies’ method. ^20^ The general form can be reduced to

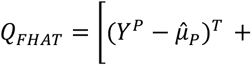

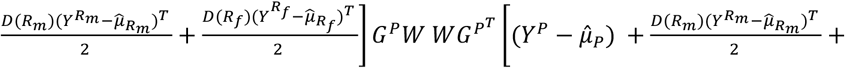

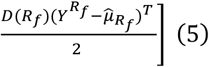

for incorporating FH from both parents (with mothers denoted by *m* and fathers denoted by *f*) when using logistic models for binary trait with the estimates of dispersion parameters fixed to 1 (i.e., 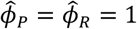, and the kinship coefficients (Ω_*m*_, Ω_*f*_) fixed to 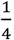.

### Optimal FHAT (FHAT-O)

Using the same framework adopted in FHAT, we developed a FHAT-O statistic based on the optimal unified test SKAT-O ^8^. Since SKAT-O combines the feature of SKAT and Burden tests, the power is robust in the presence of both different and same directions of causal variant effects. We first developed a FHAT-Burden, which is a weighted sum of the weighted score statistics in probands, and relatives based on their relationships (Supplemental Method). Then we proposed unified test defining as the weighted average of FHAT and FHAT-Burden:

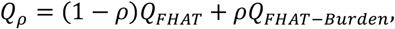

where the weight *ρ* can be estimated to minimize the p-value using the procedure proposed by Lee et al ^21^. When *ρ* = 1, *Qρ* reduces to FHAT-Burden, and when *ρ* = 0, *Qρ* is equivalent to FHAT. The statistic for optimal test FHAT-O that combines the features of FHAT and FHAT-Burden is determined as

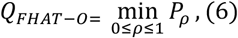

where *P*_*ρ*_ is the p-value estimated for each given *ρ* (More details are in Supplemental Method).

### Simulation Analysis

Simulations were performed to evaluate the FHAT and FHAT-O statistics in terms of empirical type I error and statistical power. We generated 10,000 haplotypes for a 4kb region on chromosome 19 using HapGen2 software ^22^. The data from 1000 genomes project was used as the reference panel to simulate haplotypes. In all simulations, we focused on binary traits because they are more often collected through questionnaire in relatives and we focused on rare variants with MAF< 1%. We used the definition from Chen et al. ^22^ to calculate the genetic effect size. We simulated the probands with both genotypes and phenotypes, and available FH data from both parents. We used LT-FH phenotype in SKAT (SKAT-LTFH) and SKAT-O (SKATO-LTFH) and compared the results to FHAT and FHAT-O, and they were all calculated by combining the FH from relatives (i.e. mothers and fathers) into the analysis. The standard methods (SKAT, SKAT-O, Burden test and ACAT-V) only used proband data. Because mothers and fathers were simulated as independent samples, they were analyzed using a single relatives’ model (3) and then FHAT and FHAT-O statistics were calculated using (5) and (6). The type I error and power of FHAT and FHAT-O were compared to SKAT-LTFH, SKATO-LTFH, SKAT, SKAT-O, Burden test and ACAT-V. Note that ACAT-V is an aggregation unit-based test combining variant-level p-values using ACAT. The detailed description of type I error and power simulation can be found in Supplemental Method.

### Analysis of Whole Exome Sequencing Data in the UK Biobank

The UK Biobank is a large prospective cohort study with information on clinical traits, covariates, and genome-wide genotype data for over 500,000 individuals with age at assessment between 37-73 years at baseline (2006 to 2010). The second tranche of exome sequence data of approximately 4 million coding variants for 200,000 individuals has been recently completed in the UK Biobank. FH of all cause dementia and hypertension was collected from questionnaires. Rare variant (with MAF< 1%) gene-based analyses detailed in the Supplemental Method were conducted to analyze all cause dementia and hypertension in the UK Biobank data.

## Results

### Type I Error and Power

A total of 2 million simulation replicates were first generated to evaluate type I error at various alpha levels for FHAT, FHAT-O, SKAT-LTFH, SKATO-LTFH, SKAT, SKAT-O, Burden test and ACAT-V using 5000 probands with available parental history (**Table 1**). When the disease prevalence is low, SKAT and SKAT-O have inflated type I error for prevalence = 20% and alpha =2.5×10^−4^ and 2.5×10^−5^, while the type I error is controlled better in FHAT, FHAT-O, SKAT-LTFH, and SKATO-LTFH when combining additional cases in relatives. When the prevalence is set to 50%, a slightly deflated type I error was observed in FHAT, SKAT-LTFH, and SKAT in some scenarios. The conservativeness of SKAT when the prevalence is 50% was also observed in prior publications ^4, 8^. The type I error evaluation results for other disease prevalence and alpha levels (including exome-wide significance) can be found in **Table S2**. We demonstrated that both FHAT and FHAT-O have reasonable type I error at the exome-wide significance (alpha= 2.5×10^− 6^) and other scenarios, and when the prevalence is low, FHAT and FHAT-O have lower inflation while other standard methods suffer from substantial inflation in type I error.

**Table 1.**
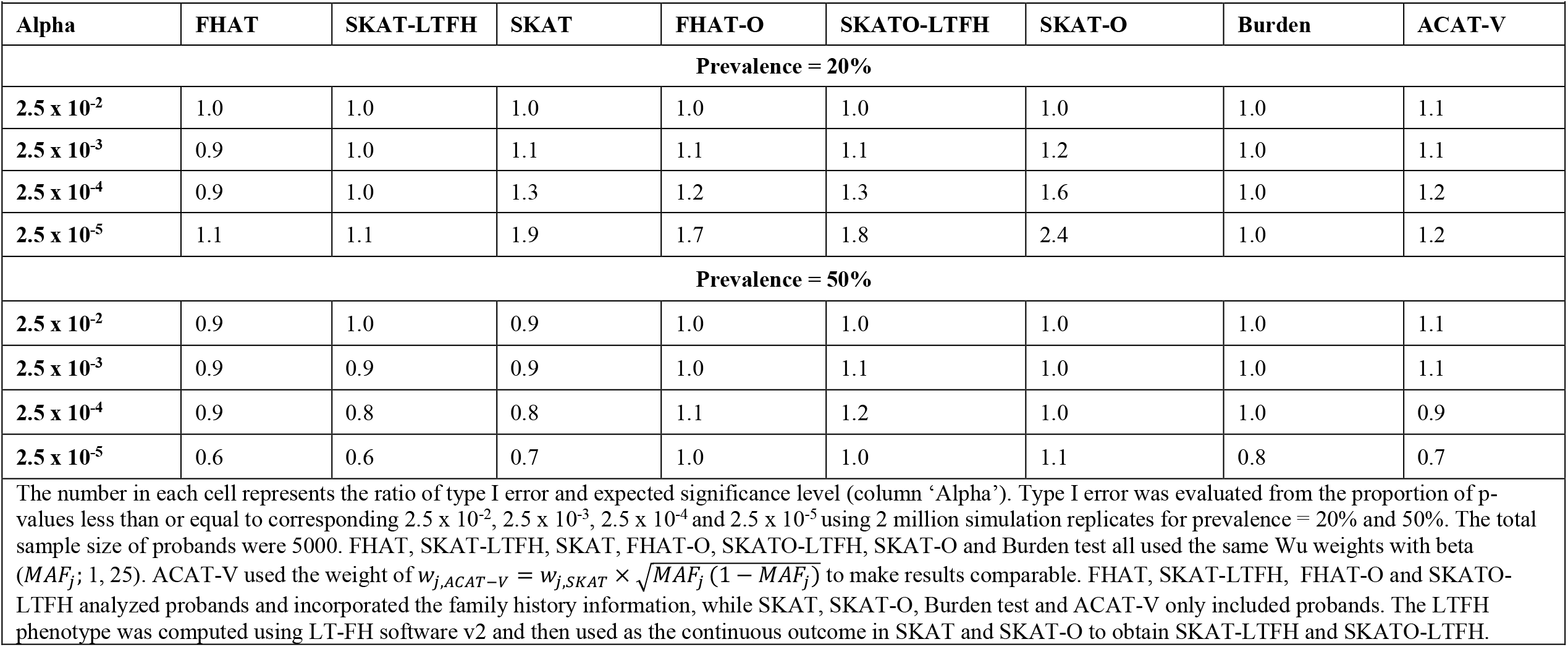
Type I Error Rates of FHAT, SKAT-LTFH, SKAT, FHAT-O, SKAT-LTFH, SKAT-O, Burden, and ACAT-V.

Figure 1 and **Figure S1** summarize the power simulation results of FHAT, SKAT-LTFH, SKAT, FHAT-O, SKATO-LTFH, SKAT-O, Burden test and ACAT-V for disease prevalence = 20% and 50%, alpha= 2.5×10^−5^ (**Figure S1**) and 2.5×10^−6^ (**Figure 1**). The causal variants in a region were set to have positive effects, or half of the causal variants have positive effects and half of the causal variants have negative effects. In all scenarios, similar patterns are shown in **Figure 1** and **Figure S1**. Our main findings included: 1) FHAT and FHAT-O are more powerful than SKAT-LTFH and SKATO-LTFH, respectively, under many scenarios when the variants have larger effects on the disease among older people; 2) FHAT and FHAT-O have greatly improved power compared to standard method that do not incorporate FH in most scenarios except for the scenario when the proportion of causal variants is 10% and half of the causal variants have positive effects and half of the causal variants have negative effects. However, ACAT-V has substantial power loss in many other scenarios; 3) FHAT suffers from a loss of power when the proportion of causal variants is high and the causal variants have effects in the same directions. In contrast, FHAT-O outperforms FHAT in those scenarios, and remains powerful regardless of the directions of genetics effects or number of causal variants.

**Figure 1.**
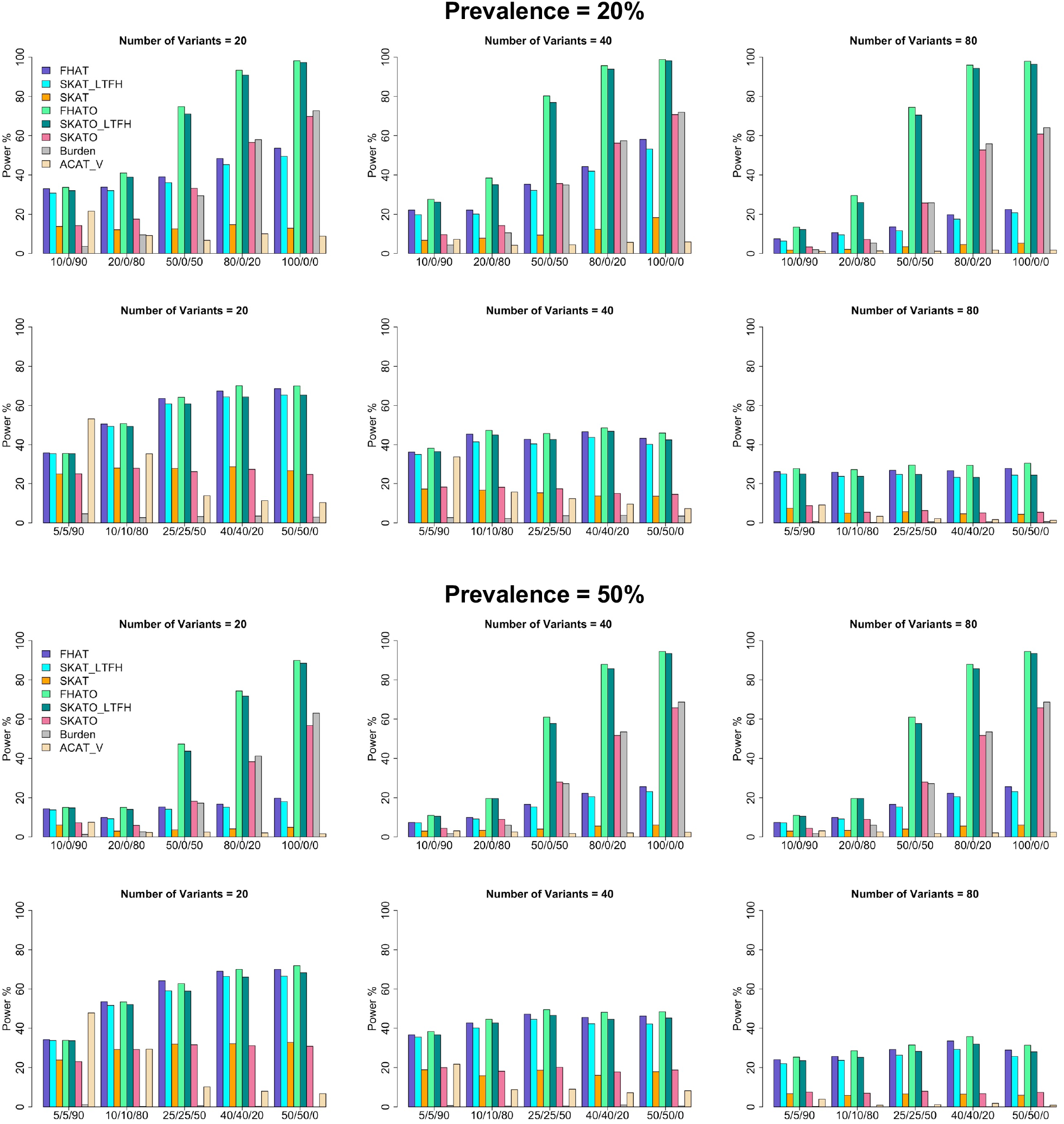
Empirical Power of FHAT, FHAT-O, SKAT-LTFH, SKATO-LTFH, SKAT, SKAT-O, Burden test and ACAT-V at Exome-wide Significance. In each plot, the x axis in the format of +/-/0 indicates the proportion of variants with positive, negative and no effects. Each bar shows the empirical power evaluated as the proportion of p-values less than or equal to alpha= 2.5 × 10^−6^. The total sample size of probands was set to 5000. The analyses were restricted to rare variants with MAF< 1%. The disease prevalence was set to 20% and 50%. FHAT, FHAT-O, SKAT-LTFH, SKATO-LTFH, SKAT, SKAT-O, and Burden test all used the same Wu weights with beta (*MAF*_*j*_; 1, 25). ACAT-V used the weight of 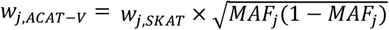 to make results comparable. FHAT, FHAT-O, SKAT-LTFH, and SKATO-LTFH analyzed probands and incorporated the family history information, while SKAT, SKAT-O, Burden test and ACAT-V only included probands. The proportion of causal variants was set to 10%, 20%, 50%, 80%, and 100%. The number of variants tested in a region considered were: 20, 40, 80.

### Computational Cost

FHAT and FHAT-O have lower computational cost compared to SKAT-LTFH and SKATO-LTFH. **Table 2** summarizes computation time (in minutes) for all methods for analyzing 1000 regions that contain 30 variants. The computation time of FHAT, FHAT-O, SKAT, SKAT-O, Burden test and ACAT-V depends on sample size and region size, whereas the running time for SKAT-LTFH and SKATO-LTFH (conducting using the LT-FH software v2 ^11^) depends on the number of configurations of probands’ disease status and FH.

**Table 2.**
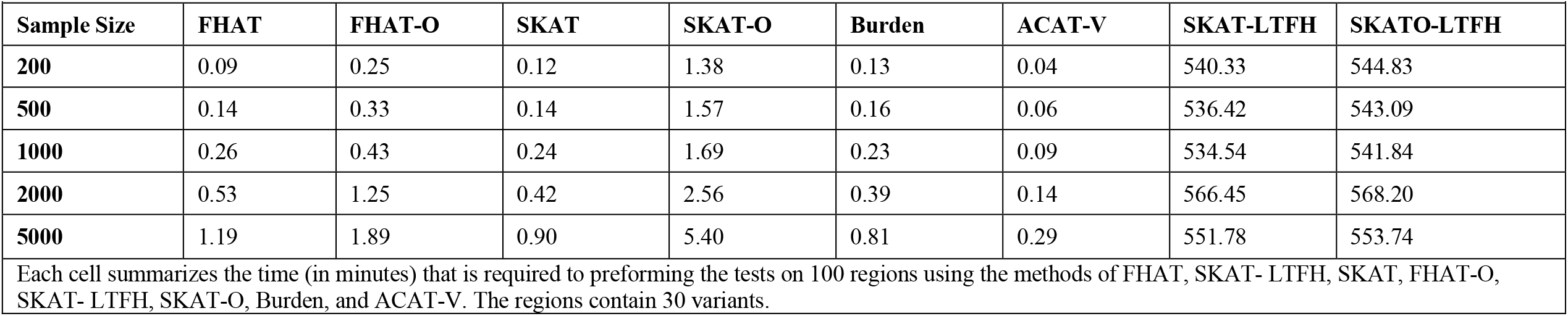
Computational Time for Testing 1000 Regions.

### Application to the UK Biobank

There are 129,670 unrelated white individuals (with ethnic background of White, British, Irish, and Any other white background) who passed all QC filters and have exome sequencing data, phenotype, and available parental disease status. The unrelated individuals were identified by removing samples who have multiple 1^st^, 2^nd^, and 3^rd^ degree relatives, randomly omitting one sample from each relative pair. The age at the first assessment visit for probands is between 38 and 72 with the mothers of probands being between 60 and 105, and the fathers of probands being between 60 and 102. There are 27 dementia cases (p=0.02%) and 32,773 hypertension cases (p=25.3%) among probands. While mothers and fathers of probands have similar hypertension prevalence (37,145 hypertension cases in mothers, p=28.6%; 26,063 hypertension cases in fathers, p= 20.1%), more dementia cases are observed in the parents (10,654 dementia cases in mothers, p=8.2%; 5,720 dementia cases in fathers, p= 4.4%) compared to probands.

We first evaluated the associations between all cause dementia and hypertension with known regions previously implicated with AD/dementia risk and hypertension. We performed the analysis for all unrelated white individuals using FHAT, FHAT-O, SKAT-LTFH, SKATO-LTFH and other conventional tests (SKAT, SKAT-O, Burden test, and ACAT-V), see results in **Table 3**. The samples involved in the analyses varied because of missing values in the covariates used for adjustment in the models. FHAT, SKAT-LTFH, FHAT-O and SKATO-LTFH had improved significance after incorporating parental phenotype information compared to p-values calculated using other conventional tests for majority of genes. SKAT, SKAT-O and ACAT-V had almost no power to detect some associations for all cause dementia due to low prevalence in probands. The results show that *BCL3* (P= 6.8×10^−5^ in FHAT, P= 2.5×10^−5^ in SKAT-LTFH, P= 5.9×10^−5^ in FHAT-O, P= 1.8×10^−5^ in SKATO-LTFH) and *TOMM4* (P= 3.0×10^−4^ in FHAT, P= 5.8 ×10^−4^ in SKAT-LTFH, P= 3.8×10^−4^ in FHAT-O, P= 7.7×10^−4^ in SKATO-LTFH) were significantly associated with all cause dementia status at a significance level of 6.3×10^−3^ for testing 8 genes. At the same significance level, *DBH* (P= 1.3×10^−3^ in FHAT, P= 2.0×10^−3^ in SKAT-LTFH, P= 2.6×10^−3^ in FHAT-O, P= 3.3×10^−3^ in SKATO-LTFH) was identified for hypertension and which had improved significance compared to the results from conventional methods. Although the tests that incorporate FH demonstrated an improved significance for all 8 AD/dementia genes we tested, some p-values for hypertension genes were less significant. This may be due to the fact that the prevalence for hypertension in probands was similar to that in parents, and the associations were diluted by the potential noises that were added when combining the FH from parents.

**Table 3.**
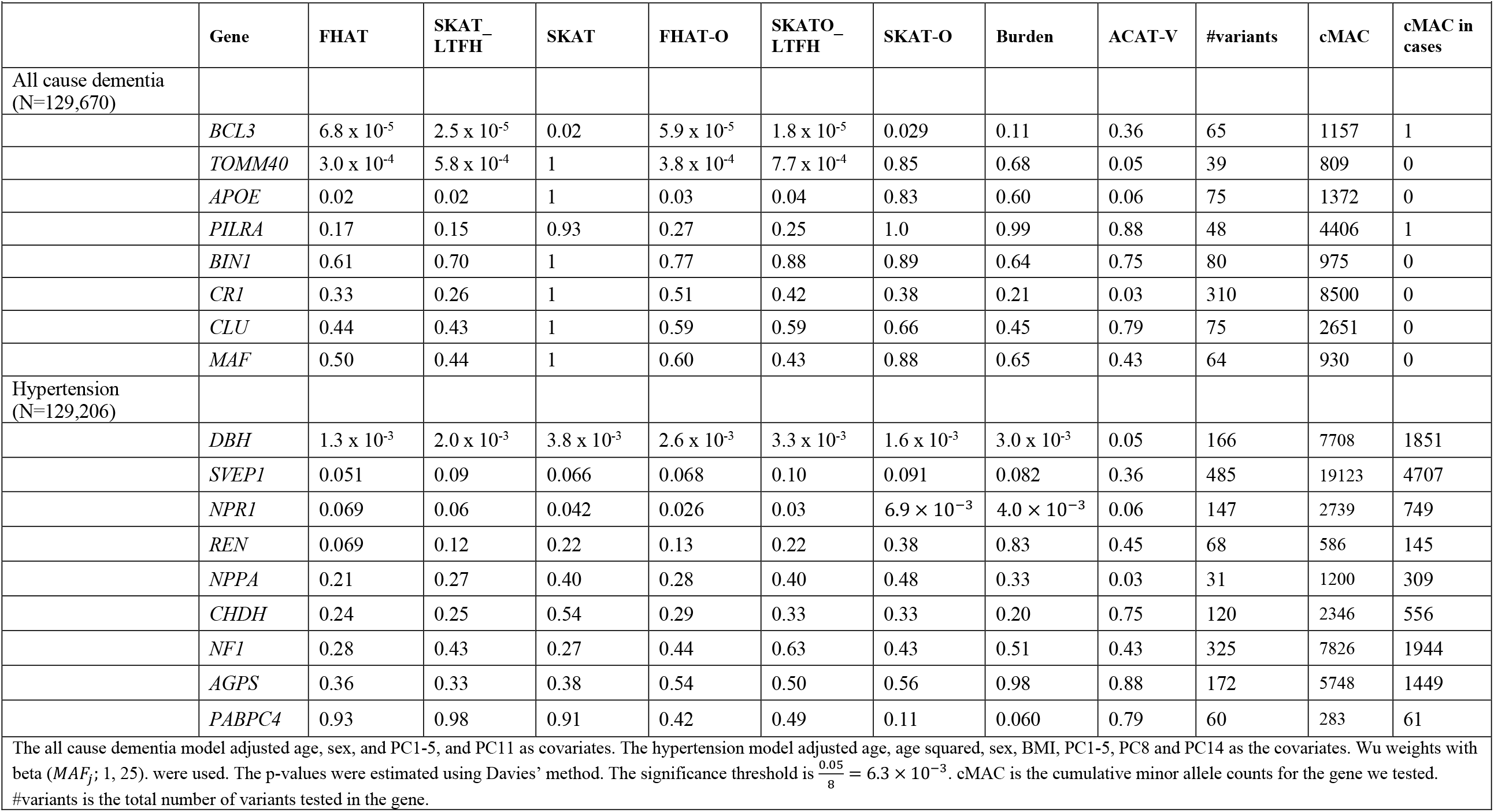
Association Analysis for Genes Previously Implicated in All Cause Dementia and Hypertension Susceptibility.

A comprehensive exome-wide analysis was then conducted. A total of ∼18K genes with two or more rare genetic variants meeting our filtering criteria were included. We used models including the same covariates for all cause dementia and hypertension as we did in the known gene analyses. In the analysis of all cause dementia (**Table 4, Figure 2**), the gene *TREM2* ^24^ (P= 4.1×10^−9^) with known effects on AD/dementia and late onset AD achieved a strict exome-wide significance (P< 2.8×10^−6^) using FHAT-O and it was also detected by FHAT (P= 5.2×10^−6^) with a suggestive exome-wide significance (P< 5.6×10^−5^). One known AD/dementia gene, *PVR* ^25^ (P= 1.2×10^−5^ in FHAT and P= 1.8×10^−5^ in FHAT-O) was identified with both FHAT and FHAT-O analysis, and *ABCA7* ^26^ (P= 4.1×10^−5^) with known effects on AD/dementia was identified by FHAT-O. Moreover, three novel genes were found to be significantly associated with all cause dementia using FHAT and FHAT-O (*EFCAB3* with P= 4.0×10^−5^ in FHAT and P= 4.2×10^−5^ in FHAT-O, *EMSY* with P= 4.4×10^−5^ in FHAT and P= 2.7×10^−5^ in FHAT-O, and *KLC3* with P= 1.4×10^−5^ in FHAT-O). Because we observed highly inflated results (**Figure 2**) from hypertension analysis due to the correlation among parents’ phenotypes, we corrected the analysis by additionally adjusting for the spouse’s hypertension status in the parents’ model. For the adjusted hypertension analysis (**Table 4, Figure 2**), FHAT identified *GATA5* (P=4.1×10^−5^), and FHAT-O identified *FGD5* (P= 4.3×10^−5^) and *DDN* (P= 4.2×10^−5^) at a suggestive significance level. Those genes detected by our methods have previously been reported to be associated with hypertension-related trait. ^27-39^

**Table 4.** Whole Exome-wide Association Analysis for All cause dementia and Hypertension.

|  | Gene name | FHAT p-value | FHAT-O p-value | #variants | cumMAC |
| --- | --- | --- | --- | --- | --- |
| **All cause dementia**<br>**(N=129,670)** |  |  |  |  |  |
| | *TREM2* | $5.2 \times 10^{-6}$ | $4.1 \times 10^{-9}$ | 45 | 4559 |
| | *PVR* | $1.2 \times 10^{-5}$ | $1.8 \times 10^{-5}$ | 75 | 2068 |
| | *EFCAB3* | $4.0 \times 10^{-5}$ | $4.2 \times 10^{-5}$ | 60 | 2579 |
| | *EMSY* | $4.4 \times 10^{-5}$ | $2.7 \times 10^{-5}$ | 158 | 1543 |
| | *KLC3* | $4.8 \times 10^{-4}$ | $1.4 \times 10^{-5}$ | 177 | 4174 |
| | *ABCA7* | $2.9 \times 10^{-3}$ | $4.1 \times 10^{-5}$ | 487 | 12179 |
| **Hypertension**<br>**(N=129,206)** |  |  |  |  |  |
| | *GATA5* | $4.1 \times 10^{-5}$ | $9.1 \times 10^{-5}$ | 88 | 5402 |
| | *FGD5* | $2.3 \times 10^{-4}$ | $4.3 \times 10^{-5}$ | 254 | 5269 |
| | *DDN* | 0.016 | $4.2 \times 10^{-5}$ | 107 | 1621 |
| The exome-wide significance threshold is $\frac{0.05}{18,000} = 2.8 \times 10^{-6}$ the suggestive exome-wide significance threshold is $\frac{1}{18,000} = 5.6 \times 10^{-5}$. cumMAC is the cumulative minor allele frequency in the region. #variants is the total number of variants in the gene. | | | | | |

**Figure 2.**
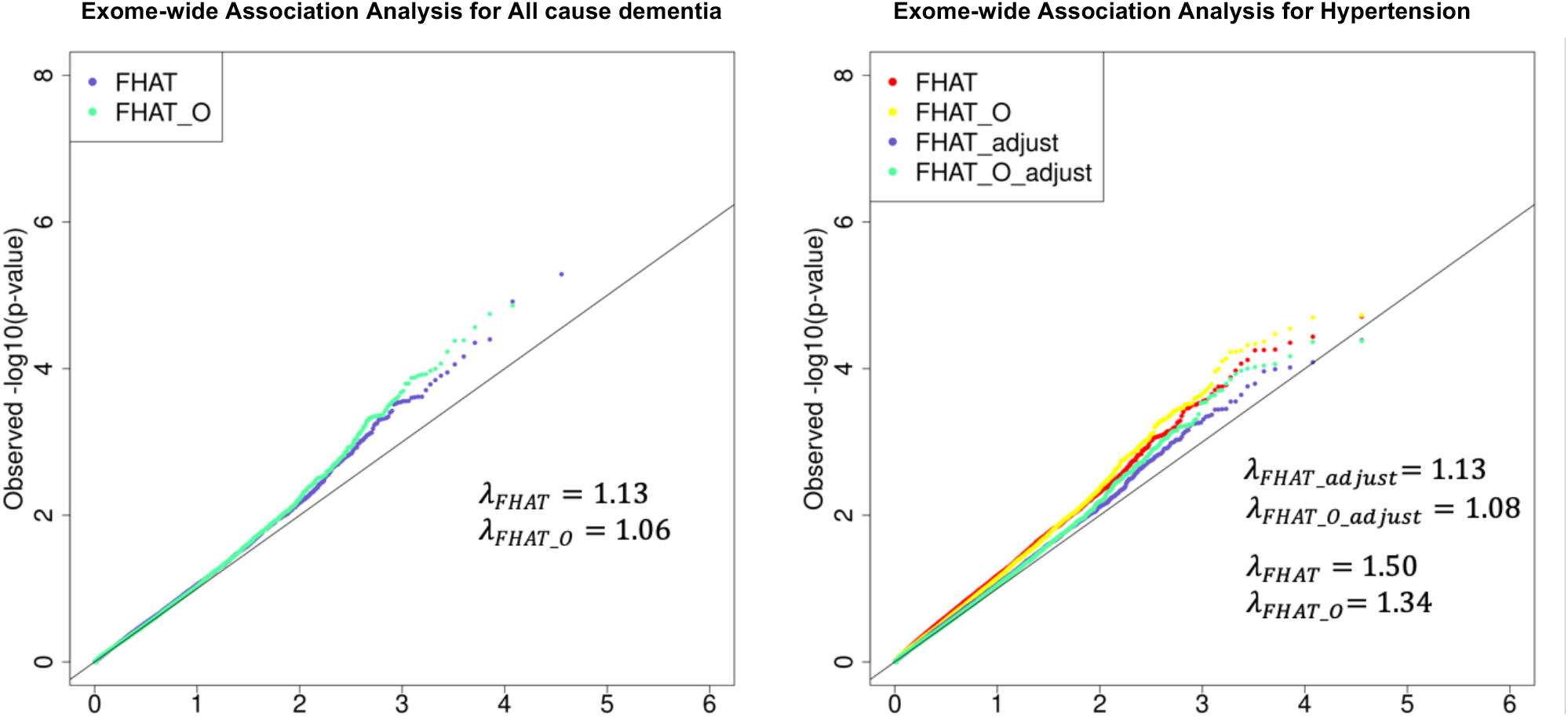
QQ Plots of Whole Exome-wide Analysis Results for All cause dementia and Hypertension. The p-values for regions with cumulative minor allele counts> 20 were used to generate the QQ plots. The left panel is the whole exome-wide analysis results for All dementia, where FHAT and FHAT-O were calculated using the model with the same covariates (age, sex, PC1-5, PC11) adjusted in AD/dementia known gene analysis. The right panel is the whole exome-wide analysis results for hypertension, where FHAT and FHAT-O were calculated using the model with the same covariates (age, age2, sex, BMI, PC1-PC5, PC8, and PC14) adjusted in hypertension known gene analysis. FHAT_adjust and FHAT-O_adjust were calculated from the adjusted hypertension analysis, where the spouse’s hypertension status combining with other previously mentioned covariates were adjusted in the parental analysis.

## Discussion

We proposed two novel approaches, FHAT and FHAT-O, that incorporate FH to increase power to detect rare variant associations in aggregation unit-based analysis. We also offered a novel way to adapt the LT-FH method to analyze rare variants. Because FH of disease is often collected through questionnaires in large cohorts, the added power is at no added cost. We applied our methods to exploit the FH from parents in simulation analysis and using the UK Biobank data, by assuming that the parents are conditionally independent. We analyzed both parents through a single relatives’ model, and combined the scores calculated from parents and probands with appropriate weights to calculate the test statistics. Because the probands’ analysis is separate from the relatives’ analysis, our methods can handle the missingness in FH as presented in (1) and (4), and one can include all probands with or without FH to optimize the usage of data.

The power was evaluated at alpha= 2.5×10^−6^ to represent the exome-wide significance for testing 20,000 genes as well as at a suggestive threshold of alpha= 2.5×10^−5^. By assuming that the causal variants in older people have bigger effects compared to younger people, we showed FHAT and FHAT-O have greater power than SKAT-LTFH, SKATO-LTFH, with greatly reduced computational cost. We also note that, as we saw a slightly higher type I error inflation in SKAT-LTFH and SKATO-LTFH than FHAT and FHAT-O, we would expect more power gain in FHAT and FHAT-O when using an empirical significance level. Compared with SKAT and ACAT-V, FHAT has greater gain in power in most cases. However, FHAT and SKAT are less powerful than Burden test and SKAT-O when there is a high proportion of causal variants, especially when the causal variants all have the same direction of effects. FHAT-O combines the features of both FHAT and FHAT-Burden, has robust power in many scenarios, and outperforms other methods, as shown in our extensive power simulations. ACAT-V has slightly higher power in some cases where the proportion of causal variants is low, which was expected because only a few genetic variants contribute to the results in ACAT-V, though the score statistic for FHAT and FHAT-O is calculated using a linear combination of squared scores from both causal and non-causal variants.

We further demonstrated that our methods have improved significance after incorporating FH from association analyses with all cause dementia and hypertension using genotypes and phenotypes collected from the UK Biobank. We compared results using FHAT, FHAT-O, SKAT-LTFH, and SKATO-LTFH for probands with both genotypes and phenotypes, and their parental history of disease to other methods only using probands. Variants in 8 known AD/dementia regions and 8 known hypertension regions were selected for the analysis. Using the significance level = 6.3×10^−3^ for testing 8 known genes, *BCL3* and *TOMM40* were significantly associated with all cause dementia while other known AD/dementia regions had improved significance compared to the methods that do not incorporate FH. Some of the hypertension genes were less significant using our method to incorporate FH, which might be caused by additional noise resulting from a similar hypertension prevalence in probands and their parents. The FHAT and FHAT-O approaches yielded similar conclusions compared to SKAT-LTFH, and SKATO-LTFH, respectively.

We evaluated type I error at various alpha levels and disease prevalence. We did not evaluate the type I error for SKAT-LTFH and SKATO-LTFH at the exome-wide significance (alpha= 2.5 × 10^−6^) to limit the computational cost. The type I error of SKAT was previously found to be conservative when the disease prevalence is ∼ 50%, and the Burden test was found to have appropriate type I error when the case-control ratio is balanced ^5-7^. However, SKAT, SKAT-O, Burden and ACAT-V suffer from substantial inflated type I error when the prevalence is low, especially for lower alpha level (i.e., alpha< 2.5×10^−4^). In contrast, the FHAT, SKAT-LTFH, FHAT-O and SKATO-LTFH control the type I error rates relatively better. The type I error is overall well controlled using FHAT and FHAT-O in most scenarios, but a high inflation occurs for alpha= 2.5×10^−6^ and prevalence = 10% where the number of cases and controls is unbalanced (**Table S2**). Unbalanced case-control ratio yields inflated type I error rates because the imbalance invalidates the asymptotic assumption of logistic regression. Saddle point approximation (SPA) ^30-32^ method and efficient resampling (ER) ^33^ have been successfully used to calibrate binary phenotype based logistic mixed models ^19^ when case-control ratios are extremely unbalanced. In the future, we plan to adopt these cutting-edge methods to properly account for unbalanced case-control ratio.

In the exome-wide association analysis, we used the same covariates (age, sex, PC1-5, PC11) as we did in the known region analysis for all cause dementia. However, as the inflation was observed in our hypertension analysis (**Figure 2**), we further adjusted for the spouse’s disease status in the parents’ model to account for the correlations among parents in addition to the covariates of age, age^2^, sex, BMI, PC1-PC5, PC8, and PC14. In the future, we will extend the current approaches to allow for correlation, as might be induced by household effect, in the analysis. Through the exome-wide analysis using FHAT and FHAT-O, we confirmed previously reported genes (*TREM2, PVR*, and *ABCA7*) ^25-27^ for AD/dementia as well as genes (*GATA5, FGD5, DDN*) ^28-30^ related to blood pressure and hypertension. Moreover, our methods identified three novel regions associated with all cause dementia (*EFCAB3, EMSY, KLC3*) using a suggestive exome-wide significance threshold. Replication analyses are needed to confirm these findings. While we observed inflated type I error for low prevalence in our simulations, we did not see evidence of large inflation of FHAT and FHAT-O in all cause dementia analysis, as seen from the Q-Q plot (**Figure 2**) and genomic control inflation factor (with *λ*_*FHAT*_ = 1.13 and *λ*_*FHAT_0*_ = 1.*06* for all cause dementia analysis).

Although the method development, simulation studies and UK Biobank analysis described in the paper were focusing on the population samples, our methods can also handle the ascertainment that happens in case-control analysis, because the likelihood can be written as the product of the retrospective proband information, taking ascertainment into consideration, and the (unascertained) relative likelihood: 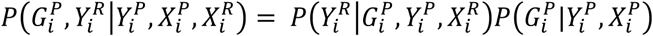 (Supplemental Method). The Equation (1) was derived based on the assumption of independence of the relatives’ phenotype and probands’ covariates conditional on the relatives’ covariate and the strength of the associations in relatives. However, when the proband covariates are believed to have an effect on the relatives’ disease status, one can adjust for such covariates in the relatives’ model (3) to account for such an effect.

In this paper, we demonstrated that FHAT and FHAT-O are computationally efficient methods compared to SKAT-LTFH and SKATO-LTFH. The significant reduced computational cost using FHAT and FHAT-O was showed in the analysis time to run 1000 aggregation unit-based tests. Although we focused on binary traits and rare variants, our method can be applied to analyze continuous traits using linear models and common variants. The framework in FHAT is flexible for various setting. While we applied FHAT and FHAT-O for probands with parental disease status available in simulations and the UK Biobank analysis, FHAT can be easily applied to other relative types. We also proposed an extension to FHAT, FHAT-O, to capture the features in SKAT-O, in particular the robustness of the power when all genetic variants have the same direction of effect and the proportion of causal variants is high. The framework can easily be extended to incorporate any other established aggregation unit-based based methods. Our methods that allow the incorporation of available FH are innovations compared to traditional rare variant studies that only use cases and controls, which have great potentials to promote genetic association research.

## Acknowledgments

Y.W., G.M.P., A.D. and J.D. acknowledge the grant support (No. 5U01AG058589) from the National Institute of Aging. H.C. acknowledge the grant support (No. R00 HL130593) from the National Institute of Health. This research was conducted using the UK Biobank Resource under Application Number 42614.

## Software

The functions for FHAT and FHAT-O are available on the website: http://sites.bu.edu/fhspl/publications/fhat/.

