## Supplemental methods, tables, and figures for "Exploiting Family History in Aggregation Unit-based Genetic Association Tests"

**Supplemental Method**

**Derivation of** $\boldsymbol{P}\left( \boldsymbol{Y}^{\boldsymbol{P}}\boldsymbol{,}\boldsymbol{Y}_{\boldsymbol{i}}^{\boldsymbol{R}} | \boldsymbol{G}_{\boldsymbol{i}}^{\boldsymbol{P}}\boldsymbol{,}\boldsymbol{X}_{\boldsymbol{i}}^{\boldsymbol{P}}\boldsymbol{,}\boldsymbol{X}_{\boldsymbol{i}}^{\boldsymbol{R}} \right)$ **and its equivalent form that takes the ascertainment into considerations**

Our methods can also account for the ascertainment in the study designs. Here we prove that the equation (1) holds and it equals to the form that enables the ascertainment adjustment in probands. In the deviations, we assume that $X_{i}^{P} \perp Y_{i}^{R}$ conditional on $X_{i}^{R}$, and $X_{i}^{R} \perp Y_{i}^{P}$ conditional on $X_{i}^{P}$.

The likelihood for the unascertained data can be written as:

$$P\left( Y^{P},Y_{i}^{R} | G_{i}^{P},X_{i}^{P},X_{i}^{R} \right)=\frac{P\left( Y^{P},Y_{i}^{R},G_{i}^{P},X_{i}^{P},X_{i}^{R} \right)}{P\left( G_{i}^{P},X_{i}^{P},X_{i}^{R} \right)}$$

$$=\frac{P\left( Y_{i}^{R} | G_{i}^{P},Y_{i}^{P},X_{i}^{R},X_{i}^{P} \right)P\left( Y_{i}^{P}|G_{i}^{P},X_{i}^{P},X_{i}^{R} \right)P\left( G_{i}^{P},X_{i}^{P},X_{i}^{R} \right)}{P\left( G_{i}^{P},X_{i}^{P},X_{i}^{R} \right)}$$

$$=P\left( Y_{i}^{R} | G_{i}^{P},Y_{i}^{P},X_{i}^{R} \right)P\left( Y_{i}^{P}|G_{i}^{P},X_{i}^{P} \right)$$

The first probability is the likelihood of family history given the proband information (and relative’s covariate information) and the second probability is the prospective likelihood of the proband status given its genotype and covariate information.

For an ascertained sample, we can write the likelihood in an alternate form:

$$P\left( G_{i}^{P},Y_{i}^{R} | Y_{i}^{P},X_{i}^{P},X_{i}^{R} \right)=\frac{P\left( Y^{P},Y_{i}^{R},G_{i}^{P},X_{i}^{P},X_{i}^{R} \right)}{P\left( Y_{i}^{P},X_{i}^{P},X_{i}^{R} \right)}=\frac{P\left( Y^{P},Y_{i}^{R} | G_{i}^{P},X_{i}^{P},X_{i}^{R} \right)P\left( G_{i}^{P},X_{i}^{P},X_{i}^{R} \right)}{P\left( Y_{i}^{P},X_{i}^{P},X_{i}^{R} \right)}$$

$$=P\left( Y_{i}^{R} | G_{i}^{P},Y_{i}^{P},X_{i}^{R} \right)\frac{P\left( Y_{i}^{P}|G_{i}^{P},X_{i}^{P} \right)P\left( G_{i}^{P},X_{i}^{P} \right)}{P\left( Y_{i}^{P},X_{i}^{P} \right)}$$

$$=P\left( Y_{i}^{R} | G_{i}^{P},Y_{i}^{P},X_{i}^{R} \right)\frac{P\left( Y_{i}^{P},G_{i}^{P},X_{i}^{P} \right)}{P\left( Y_{i}^{P},X_{i}^{P} \right)}$$

$$=P\left( Y_{i}^{R} | G_{i}^{P},Y_{i}^{P},X_{i}^{R} \right)P\left( G_{i}^{P}|Y_{i}^{P},X_{i}^{P} \right)$$

where we also assume that $G_{i}^{P} \perp X_{i}^{R}$ conditional on $X_{i}^{P}$ in addition to the assumptions for unascertained data. Again, the first probability is the likelihood of the family history given the proband information, which the second probability now refers to the standard likelihood for case-control data. Using this alternative likelihood formulation that takes ascertainment into consideration yield the same test statistics, and hence our test is valid for case-control ascertained samples.

**The null distributions of FHAT and FHAT-O**

$$Q_{FHAT}=\left[ \frac{{{(Y}^{P}-\hat{\mu}_{P})}^{T}}{\hat{\phi}_{P}}+\sum_{k} \frac{{{2\Omega_{k}D(R_{k})(Y}^{R_{k}}-\hat{\mu}_{R_{k}})}^{T}}{\hat{\phi}_{R}} \right]G^{P}W WG^{P^{T}}\left[ \frac{{{(Y}^{P}-\hat{\mu}_{P})}}{\hat{\phi}_{P}}+\sum_{k} \frac{{2\Omega_{k}D(R_{k})(Y}^{R_{k}}-\hat{\mu}_{R_{k}})}{\hat{\phi}_{R}} \right]= \sum_{j=1}^{m} \left( w_{j}^{2}S_{Pj}^{2}+\sum_{k} {4{\Omega_{k}}^{2}w}_{j}^{2}S_{{R_{k}}_{j}}^{2} \right)$$

where $w_{j}$is the pre-specified weight for variant $j$, $S_{Pj}$ is the score statistic for probands for variant $j,$ and $S_{{R_{k}}_{j}}$ is the score statistic for relative $k$ of all probands for variant $j$. Let $n$ denote the total sample size in probands, $D(R_{k})$ denote the diagonal matrix indicating the missingness of phenotypes in relative $k$ and $d_{i}(R_{k})$ denote the $i$th diagonal element from $D(R_{k})$ indicating relative $k$ for proband $i$ is missing (denoted by 0) or not (denoted by 1), then $S_{Pj}=\sum_{i=1}^{n} g_{ij}(Y_{i}^{P}-\hat{\mu}_{Pi})/\hat{\phi}_{P}$, $S_{{R_{k}}_{j}}=\sum_{i=1}^{n} {d_{i}(R_{k})g}_{ij}(Y_{i}^{R_{k}}-\hat{\mu}_{R_{k_{i}}})/\hat{\phi}_{R},$ where $g_{ij}$ is the genotype (coded as 0, 1, or 2) for the $j_{th}$ variant of $i_{th}$proband, $\hat{\mu}_{Pi}$ and $\hat{\mu}_{R_{k_{i}}}$ are the estimated mean of $Y_{i}^{P}$ and $Y_{i}^{R_{k}}$ under the null model only containing covariates, respectively, and $\hat{\phi}_{P}$ and $\hat{\phi}_{R}$ are the dispersion parameter estimates for probands and relatives, respectively.

Let $V_{P}= \left( w_{1}S_{P1},w_{2}S_{P2}, \ldots, w_{m}S_{Pm}, \right)^{T}$ and $V_{R}= \left( \sum_{k} {2{\Omega_{k}}w}_{1}S_{{R_{k}}_{1}},\sum_{k} {2{\Omega_{k}}w}_{2}S_{{R_{k}}_{2}}, \ldots, \sum_{k} {2{\Omega_{k}}w}_{m}S_{{R_{k}}_{m}}, \right)^{T}$, then $V= V_{P}+ V_{R}$ follows a multivariate normal distribution of mean 0 and covariance matrix of

$$\Psi=W{G^{P}}^{T}\left( \hat{P}+\sum_{k} 4\Omega_{k}^{2}D(R_{k})\hat{P}_{R_{k}}D(R_{k}) \right)G^{P}W,$$

where$\hat{P}={{\hat{\Sigma}^{-1}}_{P}}-{\hat{\Sigma}^{-1}}_{P}X_{P}\left( X_{P}^{T}{{\hat{\Sigma}^{-1}}_{P}}X_{P} \right)^{-1}X_{P}^{T}{\hat{\Sigma}^{-1}}_{P}$, $\hat{P}_{R_{k}}={{\hat{\Sigma}^{-1}}_{R_{k}}}-{\hat{\Sigma}^{-1}}_{R_{k}}X_{R_{k}}\left( X_{R_{k}}^{T}{{\hat{\Sigma}^{-1}}_{R_{k}}}{X_{R}}_{k} \right)^{-1}X_{R_{k}}^{T}{\hat{\Sigma}^{-1}}_{R_{k}}$ are the projection matrices in probands and relatives $k$, respectively, and ${\hat{\Sigma}_{P}}=\hat{\phi}_{P}\boldsymbol{I}$and ${\hat{\Sigma}_{R_{k}}}=\hat{\phi}_{R}\boldsymbol{I}$for continuous traits and ${\hat{\Sigma}_{P}}=diag\{1/\hat{\mu}_{Pi}(1-\hat{\mu}_{Pi})\}$and ${\hat{\Sigma}_{R_{k}}}=diag\{1/\hat{\mu}_{R_{k_{i}}}(1-\hat{\mu}_{{R_{k}}_{i}})\}$ for the binary traits. Therefore, $Q_{FHAT}= V^{T}V= \sum_{j}^{m} \lambda_{j}\chi_{1, j}^{2}$, which follows a mixture of chi-square distribution, where $\lambda_{j}$s are the eigenvalues of $\Psi$.

To test the hypothesis $H_{0}:\beta_{P}=0$, the burden score statistic derived using model (2) is

$$Q_{Burden}=\left[ \frac{1}{\hat{\phi}_{P}}\sum_{i=1}^{n} \left( Y_{i}^{P}-\hat{\mu}_{Pi} \right)\left( \sum_{j=1}^{m} w_{j}g_{ij} \right) \right]^{2}.$$

FHAT-Burden is a weighted sum of the weighted score statistics in probands, and relatives based on their relationships,

$$Q_{FHAT-Burden}=\left[ {\frac{{\sum_{i=1}^{n} \left( Y_{i}^{P}-\hat{\mu}_{Pi} \right)\left( \sum_{j=1}^{m} w_{j}g_{ij} \right)}}{\hat{\phi}_{P}}}+{\sum_{k} \frac{2{\Omega_{k}}\sum_{i=1}^{n} d_{i}(R_{k})\left( Y_{i}^{R_{k}}-\hat{\mu}_{R_{k_{i}}} \right)\left( \sum_{j=1}^{m} w_{j}g_{ij} \right)}{\hat{\phi}_{R}}} \right]^{2},$$

where $d_{i}(R_{k})$ is the $i$th diagonal element from $D(R_{k})$, indicating relative $k$ for proband $i$ is missing (denoted by 0) or not (denoted by 1).

The unified test is defined as

$$Q_{\rho}(\rho)=\left( 1-\rho\right)Q_{FHAT}+\rho Q_{FHAT-Burden}$$

$$=\left( 1-\rho\right)\sum_{j=1}^{m} (w_{j}^{2}S_{Pj}^{2}+\sum_{k} {4{\Omega_{k}}^{2}w}_{j}^{2}S_{{R_{k}}_{j}}^{2})+\rho\sum_{j=1}^{m} (w_{j}S_{Pj}+{\sum_{k} {2{\Omega_{k}}w}_{j}S_{{R_{k}}_{j}})}^{2}$$

$$={\left( 1-\rho\right)V}^{T}V+V^{T}\rho\boldsymbol{11}^{T}V= \sum_{j}^{m} \lambda_{\rho j}\chi_{1, j}^{2},$$

which asymptotically follows the distribution of a mixture-squire distribution under the null. In the summation, $\lambda_{\rho j}$s are the eigenvalues of $L_{\rho}^{T}\Psi L_{\rho}$ where $L_{\rho}$is the matrix satisfying $L_{\rho}{L_{\rho}}^{T}=\left( 1-\rho\right)\boldsymbol{I+}\rho\boldsymbol{11}^{T}$. Let $P_{\rho}$denote the p-value of $Q_{\rho}$for a given $\rho$, the test statistic for FHAT-O is

$$Q_{FHAT-O= \min_{0\leq\rho\leq1} P_{\rho},}$$

which can be evaluated from the grid search across different values of $\rho$. Defining a grid set for $\rho$as $0<\rho_{1}<\rho_{2}<\ldots<\rho_{L}$, then $Q_{FHAT-O=}\min(\rho_{1},\rho_{2},\ldots, \rho_{L})$. It can be shown that $Q_{\rho}(\rho)$ is the mixture of two independent random variables: one follows a chi-square distribution with df = 1, the other one is asymptotically approximated to the mixture of chi-square distribution with adjustment. We can use the approach proposed by Lee et al. ^21^ to obtain the optimal $\rho$.

**Liability threshold model of case–control status and family history in rare variant analysis**

The method that utilizes posterior mean generic liabilities under the liability threshold model of case-control status and FH (LT-FH) has been proposed to increase association power by combining case-control status and available FH, where they demonstrated 63% and 36% increases in power compared to GWAS and genome-wide association by proxy (GWAX) ^14^, respectively. ^11^ To make a comparison between LT-FH approach to our methods (FHAT and FHAT-O) for rare variant analysis, we first calculate posterior mean genetic liabilities Y_LT-FH_ using LT-FH conditional on both probands’ disease status and FH from relatives. Then we incorporate the LT-FH phenotype Y_LT-FH_ as a continuous outcome in SKAT and SKAT-O to test the association between aggregated groups of rare variants and the phenotype of interest.

**Simulation to Evaluate Type I Error**

Several simulation analyses were conducted under the null hypothesis of no genetic associations. The type I error was estimated using different pre-specified disease prevalence and calculated at various alpha levels. A total of 10,000 simulated haplotypes were sampled to generate 5000 probands, each assigned two haplotypes at random. Then, an unassociated phenotype $Y^{P}$ for 5000 probands and their mothers $Y^{M}$ and fathers $Y^{F}$ were simulated using the following model,

$$\binom{Y^{P}}{\begin{aligned} Y^{M} \\ Y^{F} \end{aligned}}=0.015\binom{{age}^{P}}{\begin{aligned} {age}^{M} \\ {age}^{F} \end{aligned}}+0.25\binom{{sex}^{P}}{\begin{aligned} {sex}^{M} \\ {sex}^{F} \end{aligned}}+\varepsilon, \varepsilon\sim MVN\left( 0,\Sigma\right),$$

where $age^{P},age^{M}$and $age^{F}$ are vectors of continuous variable randomly selected from ages of probands, mothers and fathers in UK biobank data, respectively; $sex^{P}$ is a vector of binary variable generated from a Bernoulli distribution with probability for female = 56% in probands;$sex^{M}$and $sex^{F}$ are fixed as female and male, respectively; $\varepsilon$ is the error term following a multivariate normal distribution with mean of zero and covariance matrix of $\Sigma$,

$$\Sigma=\delta_{g}^{2}\left( \begin{matrix} 1 & 0.5 & 0.5 \\ 0.5 & 1 & 0 \\ 0.5 & 0 & 1 \end{matrix} \right)+\delta_{e}^{2}I_{3\times3},$$

and we set $\delta_{g}^{2}=\delta_{e}^{2}=0.5$. The binary phenotype was generated using different cutoffs from the simulated continuous phenotype data to have pre-specified prevalence $p$ in probands. We fixed $p$, the prevalence in probands, at ~ 20% and 50%, with increased prevalence in mothers (~ 35% and 69%) and fathers (~ 28% and 62%), to mimic real AD/dementia data, with more prevalent disease in women and older relatives. We used $Beta \left( MAF_{j};1, 25 \right)$ weights in FHAT, FHAT-O, SKAT-LTFH, SKATO-LTFH, SKAT, and SKAT-O. The comparable weights of $w_{j, ACAT-V}=w_{j, SKAT}\times\sqrt{MAF_{j}(1-MAF_{j})}$ were used in ACAT-V. ^9^ The type I error rates were evaluated by the proportion of p-values less than or equal to the alpha levels of 2.5×10^-2^, 2.5×10^-3^, 2.5×10^-4^, and 2.5×10^-5^.

**Simulation to Evaluate Power**

The power was assessed under various scenarios. We randomly assigned two haplotypes from 10,000 simulated haplotypes to each parent, and each parent then randomly passed one of the two haplotypes without recombination to the probands. We considered a model with an interaction between age and variants to simulate a continuous phenotype

$$\binom{Y^{P}}{\begin{aligned} Y^{M} \\ Y^{F} \end{aligned}}=0.015\binom{{age}^{P}}{\begin{aligned} {age}^{M} \\ {age}^{F} \end{aligned}}+0.25\binom{{sex}^{P}}{\begin{aligned} {sex}^{M} \\ {sex}^{F} \end{aligned}}+{0.015\binom{{G_{causal}}^{P}}{\begin{aligned} {G_{causal}}^{M} \\ {G_{causal}}^{F} \end{aligned}}\gamma\binom{{age}^{P}}{\begin{aligned} {age}^{M} \\ {age}^{F} \end{aligned}}}+\varepsilon,$$

where $age$, $sex$, and $\varepsilon$ are defined in the model used for type I error simulations; $G_{causal}^{P}$,$G_{causal}^{M}$ and $G_{causal}^{F}$ are the genotype matrices of causal variants for probands, mothers and fathers in the true model, respectively, but we assumed that $G_{causal}^{M}$ and $G_{causal}^{F}$ were missing when we calculated FHAT, FHAT-O and SKAT-LTFH and SKATO-LTFH; $\gamma$ is a vector of effect sizes for the causal variants. The elements in vector $\gamma$ are specified as following, ^23^

$$\gamma_{j}=\sqrt{\frac{c}{2MAF_{j}(1-MAF_{j})},}$$

where $MAF_{j}$ is the MAF of causal variant $j$, c is a pre-specified constant and defined as

$$c=\frac{R^{2}}{V^{T}DV},$$

where $R^{2}$ is the proportion of variance explained by causal variants, we set to 2% in our simulation when all causal variants have same effect directions, and 5% when half of the causal variants have positive effects and half of the causal variants have negative effects on the liability scale for binary trait; $D$consists of the LD correlation matrix between variants; $V$ is a vector corresponding to the effect directions of causal variants. We simulated data by varying the number of total variants testing in a region, and the proportion of causal variants. The same strategy was used to generate binary phenotypes described in the type I error simulations: the binary phenotype was generated using different cutoffs for simulated continuous phenotype data.

The comparable weights were used to calculate statistics for FHAT, FHAT-O, SKAT-LTFH, SKATO-LTFH, SKAT, SKAT-O, Burden tests and ACAT-V. We considered scenarios where the number of variants analyzed in a region was 20, 40, and 80, the disease prevalence $p=$ 20% and 50%, and the proportion of causal variants is 10%, 20%, 50%, 80% and 100%. The power was calculated as the proportion of p-values less than or equal to the alpha level = 2.5×10^-5^ and exome-wide significance level = 2.5×10^-6^ for testing 20,000 genes.

**Analysis of Whole Exome Sequencing Data in the UK-Biobank**

We adjusted the all cause dementia analysis for age and sex, and adjusted the hypertension analysis for age, age squared (age^2^), sex, and body mass index (BMI). The BMI of probands was used as a proxy for BMI for the parents. To account for population structure, we additionally adjusted ancestral PCs in the analysis. The PCs from probands were used in parental analysis. We first tested the top 20 PCs in probands and parental analysis, and we meta-analyzed the results to determine which PCs to include in all cause dementia and hypertension analyses (**Table S1**). We included the top 5 PCs and any additional PCs reaching statistical significance (P< $\frac{0.05}{20}=$2.5×10^-3^) in the models: the PC1-PC5 and PC11 were included in the all cause dementia analyses, while PC1-PC5, PC8, and PC14 were included in models for hypertension. We did the variant-level quality control (QC) by removing any variants with missing rate > 5%, and then we imputed the missing genotypes using the mean genotype values, as is standard in the SKAT method. The following QC procedures were implemented to select samples: samples with high heterozygosity and high missing rates, sex aneuploidy, and mismatches reported sex with affymetrix-determined sex were removed from our analysis. We selected frameshift, splice accepter, splice donor, stop gained, stop lost, and missense variants determined by Ensemble Variant Effect Predictor (VEP) annotation for inclusion in our analyses. We used the genotypes generated from Functionally Equivalent (FE) pipeline and we omitted regions flagged as unreliable in the UK Biobank exome sequencing data.

**Supplemental Tables**

**Association analysis between PCs and Disease of Interest**

Before selecting the final model for all cause dementia and hypertension analysis, we first tested the significance between diseases (all cause dementia and hypertension) and each of the 20 PCs in parents and probands separately, and meta-analyzed the results. We used the probands’ PCs as the proxy-PCs for relatives. We selected the top 5 PCs and significant PCs (P< $\frac{0.05}{20}=$2.5×10^-3^) to adjust for population structure in the analysis.

| **Table S1. P-values for the Association Analysis Between PCs and Diseases** | | |
| --- | --- | --- |
|  | All cause dementia | Hypertension |
| PC1 | 0.01 | 0.71 |
| PC2 | 0.53 | 0.37 |
| PC3 | 0.51 | 0.4 |
| PC4 | 0.31 | 4.0 x 10^-3^ |
| PC5 | 2.0 x 10^-3^ | 0.024 |
| PC6 | 0.96 | 0.02 |
| PC7 | 0.26 | 0.11 |
| PC8 | 0.037 | 9.3 x 10^-5^ |
| PC9 | 0.19 | 0.045 |
| PC10 | 0.27 | 0.89 |
| PC11 | 6.2 x 10^-4^ | 0.16 |
| PC12 | 0.56 | 0.11 |
| PC13 | 0.34 | 0.53 |
| PC14 | 0.2 | 6.8 x 10^-6^ |
| PC15 | 0.78 | 0.14 |
| PC16 | 6.6 x 10^-3^ | 0.052 |
| PC17 | 0.028 | 0.2 |
| PC18 | 0.24 | 0.81 |
| PC19 | 0.78 | 0.82 |
| PC20 | 0.16 | 3.7 x 10^-3^ |
| The top 5 PCs and significant PCs were selected in the testing models. Using the significance threshold of 2.5 x 10^-3^ for testing 20 PCs, we adjusted PC1-5, and PC11 were adjusted in all cause dementia model, and PC1-5, PC8 and PC14 were adjusted in hypertension model. | | |

**Simulations for Type I error at exome-wide significance and other disease prevalence**

The Type I error for FHAT and FHAT-O was evaluated at exome-wide significance and compared to other methods. Here we did not compare to SKAT-LTFH and SKATO-LTFH due to a high CUP time consumption. We considered the disease prevalence = 10%, 20% 30% and 50%, and we generated 20 million replicates for each scenario. With $p$being fixed as ~ 10%, 20%, 30% and 50%, we had the prevalence ~ 21%, 35%, 41%, and 69% in mothers, and the prevalence ~ 16%, 28%, 49%, and 62% in fathers. We first simulated $Y^{P}$ for 5000 probands and their mothers $Y^{M}$ and fathers $Y^{F}$ using the same simulation model presented in the Simulation to Evaluate Type I Error section.

| **Table S2. Type I Error Rates of FHAT, FHAT-O, SKAT, SKAT-O, Burden, and ACAT-V** | | | | | | |
| --- | --- | --- | --- | --- | --- | --- |
| **Prevalence** | **FHAT** | **SKAT** | **FHAT-O** | **SKAT-O** | **Burden** | **ACAT-V** |
| **Alpha= 2.5 x 10^-4^** | | | | | | |
| **P=10%** | 1.4 | 2.4 | 1.6 | 2.7 | 1.2 | 1.3 |
| **P=20%** | 0.9 | 1.3 | 1.2 | 1.6 | 1.0 | 1.2 |
| **P=30%** | 0.9 | 1.0 | 1.2 | 1.2 | 1.0 | 1.1 |
| **P=50%** | 0.9 | 0.8 | 1.1 | 1.0 | 1.0 | 0.9 |
| **Alpha= 2.5 x 10^-6^** | | | | | | |
| **P=10%** | 3.2 | 8.6 | 3.8 | 10.9 | 2.0 | 2.1 |
| **P=20%** | 1.3 | 2.7 | 2.0 | 4.0 | 1.6 | 1.2 |
| **P=30%** | 0.8 | 1.5 | 1.4 | 2.4 | 1.3 | 1.3 |
| **P=50%** | 0.6 | 0.5 | 1.0 | 1.0 | 1.0 | 0.9 |
| The number in each cell represents the ratio of type I error and expected significance level. Type I error was evaluated from the proportion of p-values less than or equal to corresponding 2.5 x 10^-4^ and 2.5 x 10^-6^ using 20 million simulation replicates. The x axis is the disease prevalence of 5%, 10%, and 20%. The total sample size of probands were 5000. FHAT FHAT-O, SKAT, SKAT-O and Burden test all used the same Wu weights with beta ($MAF_{j};$1, 25). ACAT-V used the weight of $w_{j, ACAT-V}=w_{j, SKAT}\times\sqrt{MAF_{j}(1-MAF_{j})}$ to make results comparable. FHAT and FHAT-O analyzed probands and incorporated the family history information, while SKAT, SKAT-O, Burden test and ACAT-V only included probands. | | | | | | |

**Supplemental Figures**

**Empirical power at Alpha= 2.5 x 10^-5^**

**Prevalence = 20%**

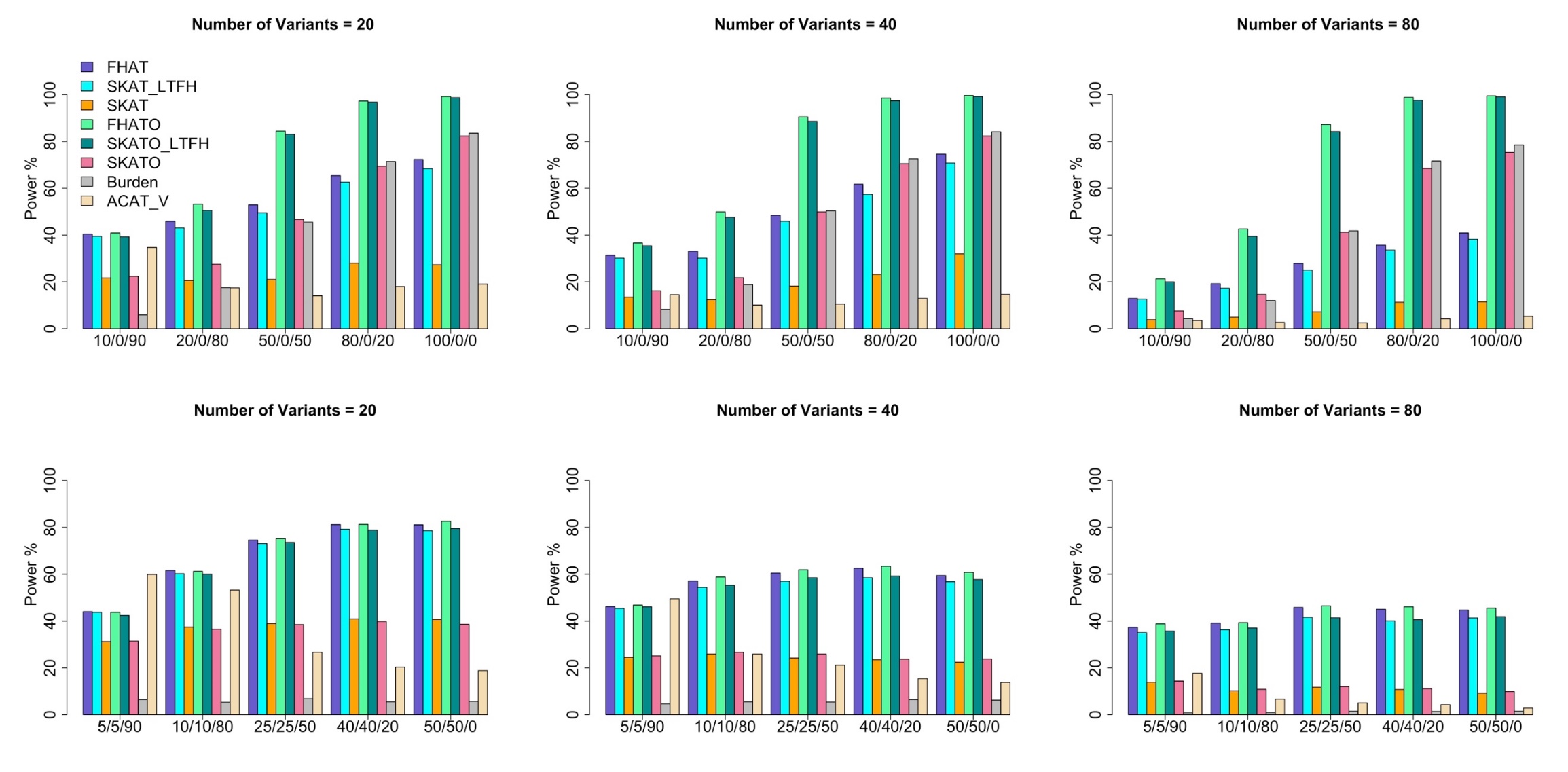

**Prevalence = 50%**

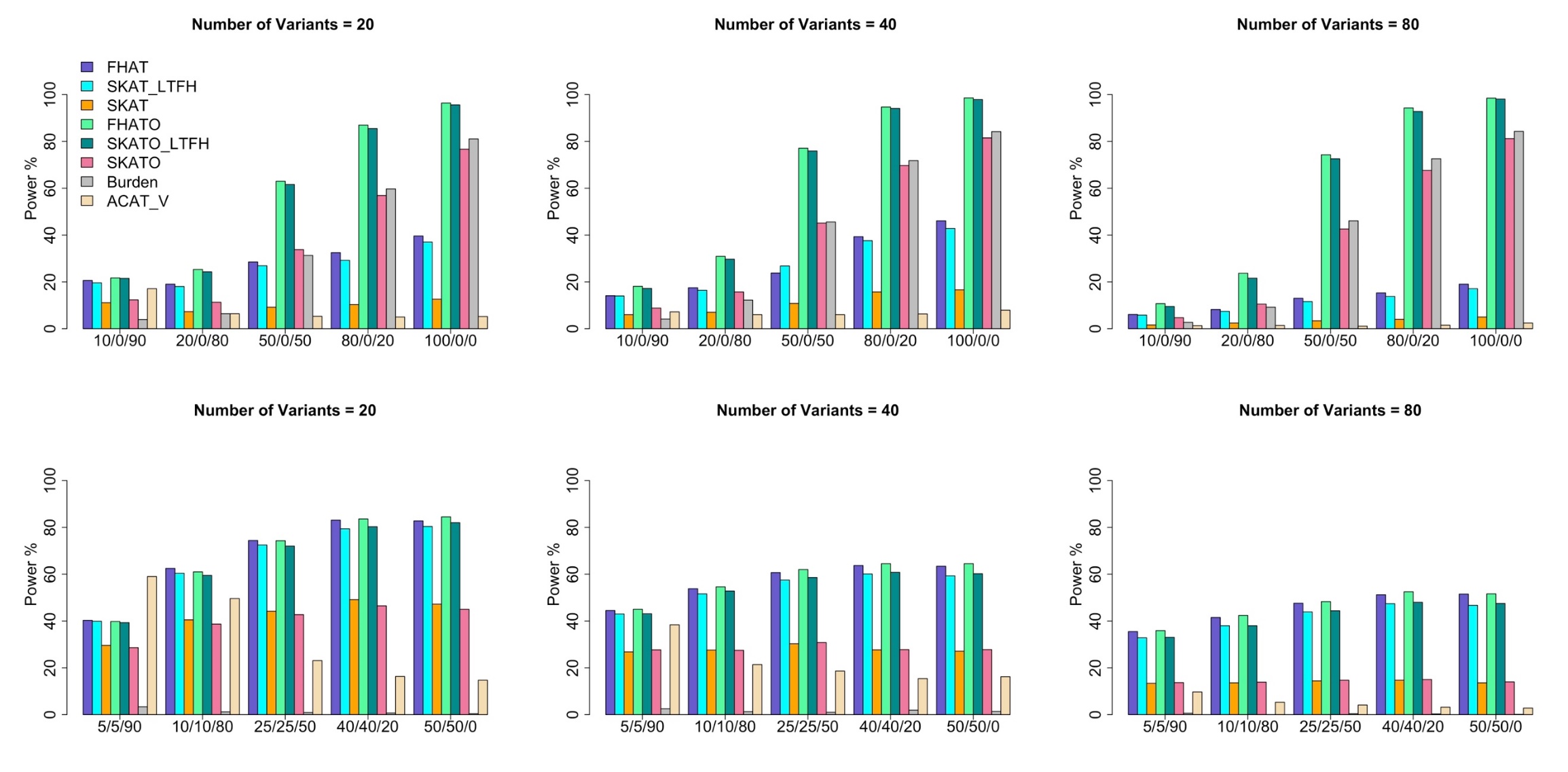

**Figure S1. Empirical Power of FHAT, FHAT-O, SKAT-LTFH, SKATO-LTFH, SKAT, SKAT-O, Burden test and ACAT-V at Alpha= 2.5 x 10^-5^**

In each plot, the x axis in the format of +/-/0 indicates the proportion of variants with positive, negative and no effects. Each bar shows the empirical power evaluated as the proportion of p-values less than or equal to alpha= 2.5 x 10^-5^. The total sample size of probands was set to 5000. The analyses were restricted to rare variants with MAF< 1%. The disease prevalence was set to 20% and 50%. FHAT, FHAT-O, SKAT-LTFH, SKATO-LTFH, SKAT, SKAT-O, and Burden test all used the same Wu weights with beta ($MAF_{j};$1, 25). ACAT-V used the weight of $w_{j, ACAT-V}=w_{j, SKAT}\times\sqrt{MAF_{j}(1-MAF_{j})}$ to make results comparable. FHAT, FHAT-O, SKAT-LTFH, and SKATO-LTFH analyzed probands and incorporated the family history information, while SKAT, SKAT-O, Burden test and ACAT-V only included probands. The proportion of causal variants was set to 10%, 20%, 50%, 80%, and 100%. The number of variants tested in a region considered were: 20, 40, 80.
